# Stage-specific checkpoint signalling by ATM, ATR and DNA-PK safeguards oocyte genome integrity

**DOI:** 10.64898/2026.10.13.758765

**Authors:** Georgios Sioutas, Madeline J. Perry, Agata P. Zielinska, Elias Elinati, Todd Fallesen, Vanessa Dao, Obah Ojarikre, James M. A. Turner

## Abstract

In somatic cells, DNA double-strand breaks (DSBs) are potentially catastrophic lesions that activate phosphatidylinositol 3-kinase-related kinase (PIKK) signalling to promote DNA repair or eliminate damaged cells. During meiosis, however, DSBs are deliberately generated as essential intermediates in homologous recombination, raising the question of how DNA damage signalling distinguishes programmed recombination from genome instability. To define how PIKK signalling is adapted to these distinct contexts, we resolved the function of the three PIKKs ATM, ATR and DNA-PK in physiological meiotic recombination and the response to postnatal DNA damage in mouse oocytes. Following postnatal irradiation, ATM functions as the dominant response kinase, with ATR and DNA-PK acting redundantly to eliminate damaged oocytes. In contrast, meiotic surveillance is mediated by ATR, with ATR removal rescuing oocyte loss in response to asynapsis and, independently, to persistent meiotic DSBs. Our findings demonstrate that DNA damage signalling is developmentally reconfigured during oogenesis and highlight distinct pathways of oocyte elimination that may ultimately inform strategies to maintain the ovarian reserve in mammals.

## Introduction

Oocytes of most mammalian species are formed during embryonic development, resulting in a finite number of eggs in adult females that lie dormant in primordial follicles.^1^ The size of this oocyte pool, termed the ovarian reserve, as well as their quality, will determine an individual’s reproductive lifespan. Oocytes are produced through the process of meiosis, the specialised cell division that results in the generation of haploid gametes from diploid germ cells. The importance of healthy oocytes for reproductive success necessitates the existence of checkpoints, which together ensure that compromised oocytes are eliminated. Despite their significance, how these checkpoints function at the molecular level is incompletely understood.

Meiotic prophase I begins with programmed double-strand breaks (DSBs), catalysed by the enzyme SPO11 during leptonema.^2–4^ DSBs are then resected, forming single-stranded DNA tails that invade the uncut homologous template, facilitated by recombinases DMC1 and RAD51.^5^ This invasion drives homologous synapsis, which initiates in zygonema with the formation of the synaptonemal complex, and complete synapsis marks the entry to pachynema. At this stage, a subset of recombination intermediates is ultimately resolved as crossovers, the physical attachments essential for homolog biorientation at metaphase I.^6^ Following crossover formation, cells progress into diplonema with the desynapsis of homologous chromosomes, after which oocytes arrest at the dictyate stage.

Previous studies have demonstrated the existence of three distinct checkpoint responses that eliminate defective oocytes: the synapsis checkpoint, the meiotic recombination checkpoint, and the postnatal DNA damage checkpoint.^7–11^ The first two operate during foetal life, removing oocytes with homologous asynapsis or unrepaired recombination intermediates. The third operates after birth, eliminating dictyate-arrested oocytes exposed to exogenous DNA damaging agents, e.g., radiation.

The synapsis, meiotic recombination and postnatal DNA damage checkpoints exhibit overlapping and distinct properties. The synapsis checkpoint requires the HORMA-domain proteins HORMAD1 and HORMAD2: typically displaced at the beginning of pachynema, prolonged retention of HORMAD1/2 on meiotic axes signals asynapsis.^12–17^ Additionally, the tumour suppressor protein BRCA1 is recruited to asynapsed chromosomes in a HORMAD1/2-dependent manner and is important for efficient checkpoint activation.^18^ In contrast, the recombination checkpoint is less well-understood. While it has been established that the SUMO ligase RNF212 promotes the detection of unrepaired recombination intermediates by blocking inter-sister repair on desynapsing axes,^19^ how these intermediates are monitored during pachynema remains unresolved. Both the synapsis and recombination checkpoints are thought to activate checkpoint kinases CHK1 and CHK2,^20^ while the postnatal DNA damage checkpoint depends largely on CHK2.^7^

A major question in oocyte checkpoint biology concerns the identity of the apical kinases that activate these three checkpoints. Primary candidates are the PIKK family members ATM, ATR and DNA-PK. ATR localises to asynapsed chromosome axes and unresolved meiotic DSBs,^12,15,21,22^ positioning it as a candidate for meiotic surveillance. However, studies in male mice have also implicated ATM in pachytene checkpoint control.^23^ ATM additionally regulates formation and processing of programmed DSBs,^24^ complicating efforts to distinguish its roles in normal meiosis from those in checkpoint activation. By contrast, less is known about the role of ATR and DNA-PK in female meiotic checkpoint control. Addressing these questions requires temporally controlled deletion that separates PIKK functions in unperturbed meiosis from checkpoint activation.

Here, we use temporally resolved, combinatorial deletions to systematically examine the functions of the PIKK network across all three oocyte checkpoints. We find that surveillance of synapsis and meiotic recombination is controlled by ATR. However, after birth, the PIKK hierarchy is reconfigured, with ATM becoming the dominant kinase controlling elimination of oocytes with exogenous damage. We propose that this developmental reorganisation reflects the specialised functions of ATM during meiotic prophase, which constrain its deployment as a canonical checkpoint kinase while programmed recombination is underway.

## Results

### The postnatal oocyte DNA damage response to irradiation centres on ATM, with redundant contributions by ATR and DNA-PK

Postnatal oocytes in primordial follicles are highly sensitive to irradiation-induced DNA damage, and low doses of γ-radiation result in complete oocyte loss.^7^ To determine the role of PIKKs in response to irradiation, we combined conditional loxP (floxed, fl) alleles of *Atm* and *Atr* with a *Ddx4-Cre* transgene which drives Cre expression from embryonic day 15.5 (E15.5).^25^ We observed detectable protein loss from birth (hereafter termed postnatal day 0; P0) (**Figure S1A, B**). This timing allowed us to distinguish between the roles of ATM and ATR during prophase I and their checkpoint role in neonatal oocytes. We irradiated mice at P5 and assessed oocyte survival at P10, by immunostaining with the oocyte-specific marker MVH. To examine the role of DNA-PK, we used an established constitutive allele for its catalytic subunit (*Prkdc^scid^*).

Individual depletion of ATM (*Atm^fl/-^Ddx4-Cre*), ATR (*Atr^fl/-^ Ddx4-Cre*) or DNA-PK (*Prkdc^scid/scid^*) did not rescue oocyte loss: ovaries were indistinguishable from irradiated controls (**Figure S1C, D**). This finding raised the possibility of functional overlap between PIKKs in regulating oocyte elimination in response to DNA damage. To examine this possibility, we generated combinatorial deletions of ATM, ATR and DNA-PK. Combined depletion of ATM and DNA-PK, or ATM and ATR, partially rescued primordial follicle elimination. However, co-depletion of ATR and DNA-PK did not (**Figure 1A, B**). This result demonstrated that ATM is crucial for oocyte elimination after irradiation, with redundant contributions by ATR and DNA-PK. Importantly, co-deletion of all PIKKs (*Atm^fl/-^ Atr^fl/-^ Prkdc^scid/scid^ Ddx4-Cre*, hereafter termed triple KO: TKO) completely rescued the oocyte loss phenotype (**Figure 1A, B**). This rescue was not transient, as irradiated TKO ovaries retained oocytes at P21 (**Figure S1E, F**). We conclude that oocyte elimination after irradiation requires PIKK-dependent DNA damage signalling, with ATM as the principal kinase.

**Figure 1.**
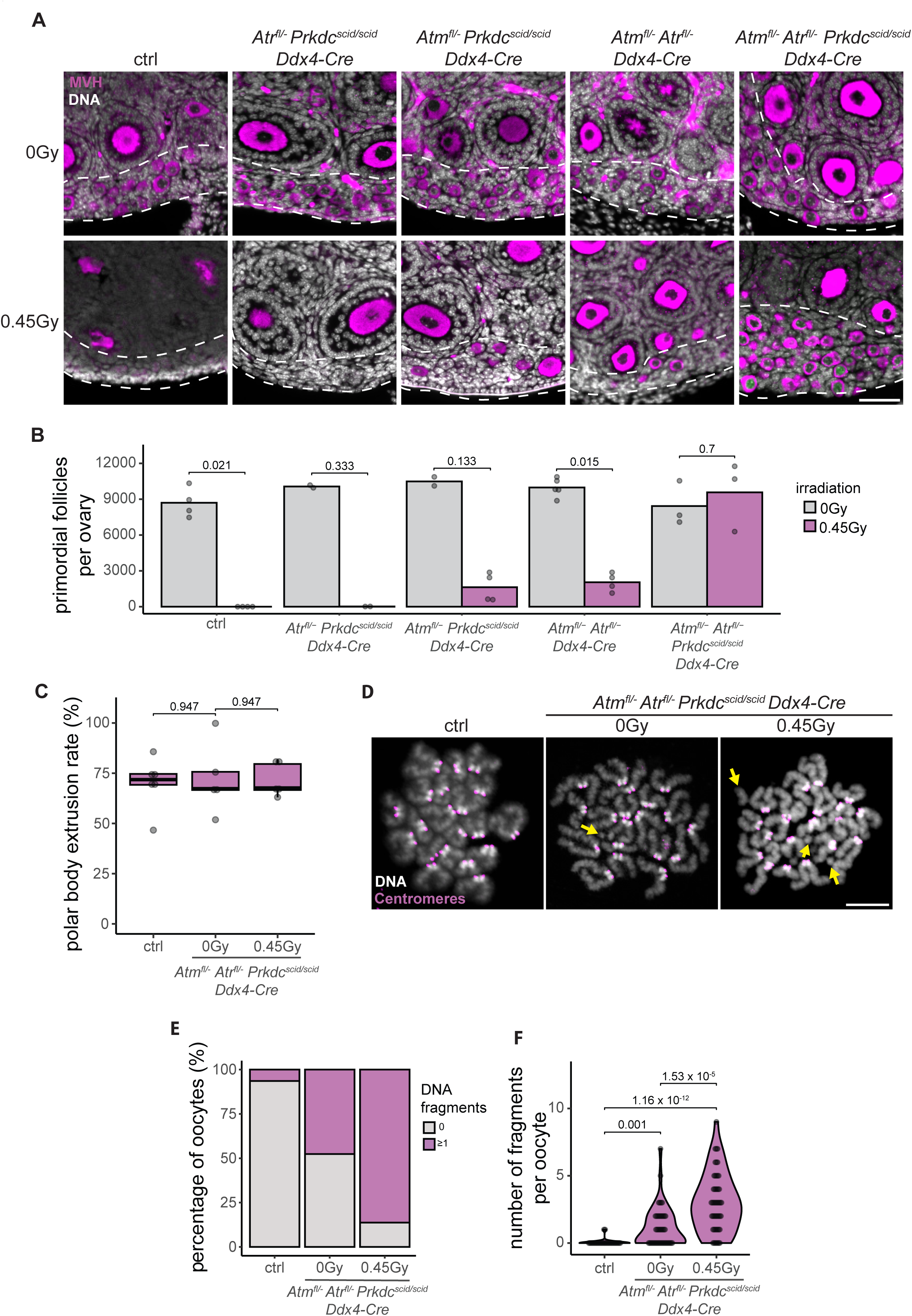
ATM, ATR and DNA-PK are required for the postnatal oocyte DNA damage response. **A)** Widefield images of sections of P10 ovaries from control, *Atr^fl/-^ Prkdc^scid/scid^ Ddx4- Cre*, *Atm^fl/-^ Prkdc^scid/scid^ Ddx4-Cre*, *Atm^fl/-^ Atr^fl/-^ Ddx4-Cre* and *Atm^fl/-^ Atr^fl/-^ Prkdc^scid/scid^ Ddx4-Cre* mice, without (0Gy) or after treatment with 0.45Gy of γ-radiation at P5, immunostained for MVH (magenta) and counterstained with DAPI (DNA, grey). Dashed white lines indicate the ovarian cortex, where primordial follicles are located. Scale bar indicates 50μm. **B)** Bar plot showing the average number of primordial follicles in all genotypes from (**A**), without irradiation treatment (0Gy) and after 0.45Gy of irradiation treatment at P5. Bar height represents average primordial follicle numbers. Dots represent follicle counts from individual animals. Number of females used: 4 (0Gy) and 4 (0.45Gy) for control, 2 (0Gy) and 2 (0.45Gy) for *Atr^fl/-^ Prkdc^scid/scid^ Ddx4-Cre*, 2 (0Gy) and 4 (0.45Gy) for *Atm^fl/-^ Prkdc^scid/scid^ Ddx4-Cre*, 5 (0Gy) and 4 (0.45Gy) for *Atm^fl/-^ Atr^fl/-^ Ddx4-Cre* and 3 (0Gy) and 3 (0.45Gy) for *Atm^fl/-^ Atr^fl/-^ Prkdc^scid/scid^ Ddx4-Cre*. p values were calculated using Mann Whitney U test. **C)** Box and whiskers plot of polar body extrusion rates from control, and *Atm^fl/-^ Atr^fl/-^ Prkdc^scid/scid^ Ddx4-Cre* mice without (0Gy) or after 0.45Gy irradiation treatment at P5. Dots represent the percentage of GV-oocytes from a single animal that extruded a polar body. 183 (from 6 females), 193 (from 5 females) and 181 (from 5 females) oocytes were assessed, respectively. **D)** Chromosome spreads of metaphase II oocytes from the same genotypes and conditions as (**C**), stained using CREST serum (centromeres, magenta) and counterstained with Hoechst 33342 (DNA, greys). Yellow arrows indicate chromosome fragments. Scale bar represents 10μm. **E)** Stacked bar plot showing the percentage of oocytes from mice of the same genotype and conditions as (**c**), that contained no fragment, or at least one fragment at metaphase II. **F**) Violin plot showing the number of fragments detected in the same oocyte pool as (**E**). Dots represent individual oocytes. 31 (from 2 females), 42 (from 4 females) and 51 (from 4 females) oocytes were assessed, respectively. p values were calculated using the Kruskal-Wallis test, followed by Dunn’s post hoc test with Benjamini-Hochberg correction.

### Rescued oocytes resume meiosis but harbour chromosome damage

Oocytes of primordial follicles lie dormant in the dictyate stage of prophase I until they are activated to grow into mature oocytes. Fully grown germinal vesicle (GV) oocytes resume meiosis upon recruitment for ovulation, with successful meiotic resumption marked by extrusion of the first polar body.^26^ To determine whether rescued oocytes retain the capacity to resume meiosis, we assessed polar body extrusion following irradiation. Non-irradiated and irradiated TKO oocytes resumed meiosis at similar rates to control oocytes, demonstrating that neither PIKK deletion nor irradiation substantially impair meiotic resumption (**Figure 1C**).

We next asked whether rescued oocytes retained chromosome integrity. Examination of metaphase II chromosome spreads revealed very low levels of chromosome fragmentation in control oocytes (6.45%). However, more than 80% of irradiated TKO oocytes contained at least one chromosome fragment (**Figure 1D, E**). Unexpectedly, approximately 50% of non-irradiated TKO oocytes also contained chromosome fragments, indicating that PIKK-depleted oocytes exhibit excessive chromosome damage even in the absence of an exogenous damage source. The chromosome damage phenotype was also evident when quantified by the number of fragments per oocyte. Non-irradiated TKO oocytes contained more fragments than controls, while irradiation treatment resulted in even more chromosome fragments in TKO oocytes (**Figure 1F**).

To determine which PIKK was required for preventing this chromosome fragmentation phenotype, we examined metaphase II spreads from non-irradiated mice carrying individual and combinatorial PIKK mutations. Oocytes from all genotypes examined resumed meiosis at similar rates (**Figure S1H**). Chromosome fragmentation was minimal in oocytes lacking DNA-PK, ATM or both DNA-PK and ATR, but increased to approximately 30% of oocytes when ATM depletion was combined with either DNA-PK or ATR depletion. Therefore, even though ATM plays a prominent role in DSB repair in postnatal oocytes, fragmentation is seen in 50% of oocytes only when all three PIKKs are removed (**Figure S1G, I, J**). The lesions observed could represent residual meiotic DSBs, or *de novo* DSBs arising after birth that are normally repaired by PIKKs. While we cannot distinguish between these possibilities, we can conclude that redundant PIKK signalling is required to maintain chromosome integrity in postnatal oocytes.

### Maternal ATR is required for preimplantation development

We next asked whether rescued oocytes were competent to support fertilisation and embryonic development. Non-irradiated and irradiated TKO females were mated to wild-type males. Oocytes from both groups could be successfully fertilised and formed zygotes, demonstrating that neither PIKK depletion nor irradiation prevents fertilisation (**Figure S2A, B**). However, embryos derived from non-irradiated and irradiated TKO mothers exhibited severe defects in preimplantation development. In both cases, embryos did not develop beyond the 4-cell stage (**Figure S2A, B**). The finding that all embryos from non-irradiated TKO females failed to develop suggested that the phenotype could not be solely explained by unrepaired DSBs, which were evident only in 50% of oocytes (**Figure 1E**). To identify the PIKK responsible for this maternal-effect phenotype, we assessed embryonic development in females carrying individual and combinatorial PIKK mutations. Approximately 60% of embryos derived from mothers lacking ATM and DNA-PK (i.e. only expressed ATR) developed to the morula stage (**Figure S2C, D**), consistent with a high percentage of oocytes containing chromosome fragments (**Figure S1I**). In contrast, loss of ATR alone was sufficient to impair preimplantation development, with most embryos failing to progress beyond the 4-cell stage (**Figure S2C, D**). Thus, the developmental failure of embryos derived from PIKK-depleted oocytes is associated with maternal ATR deficiency, in addition to the chromosome fragmentation phenotype caused by PIKK removal. These findings identify ATR as a maternal-effect protein required for normal preimplantation development.

### Early depletion of ATR does not disrupt meiotic progression

Having established a central role for ATM in the postnatal DNA damage checkpoint, we next assessed which of the PIKKs regulates the synapsis and recombination checkpoints. We began by assessing the role of ATR in unperturbed female meiosis, which to date has only been studied using a hypomorphic allele.^27^ To achieve early meiotic ATR depletion, we utilised the *Stra8-GFPCre* allele, which is expressed around meiotic entry, from E12.5.

ATR protein first became detectable in control embryonic ovaries at E15.5. ATR levels were lower in mutant ovaries of the same age, confirming successful Cre-mediated depletion (**Figure S3A**). In contrast to the mid pachynema arrest observed in male *Atr* mutants,^21,22^ ATR depletion in *Atr^fl/-^ Stra8-GFPCre* females did not disrupt meiotic progression. We assessed DSB processing dynamics by immunostaining for RPA2, which marks the single-stranded DNA intermediates of active DSB processing.^28^ Pachytene oocytes at both E16.5 and P0 contained comparable numbers of RPA2 foci in control and mutant ovaries, suggesting that DSB processing was grossly unaffected (**Figure S3B, C**). Furthermore, the incidence of asynapsis, assayed by persistent HORMAD2 staining in pachytene oocytes from E18.5 ovaries, was comparable to control (**Figure S3D, E**). Ovaries from *Atr^fl/-^ Stra8-GFPCre* mice consistently showed an elevated number of primordial follicles at P21 when compared to controls, although this did not reach statistical significance (**Figure S3F, G**). We conclude that early ATR depletion does not affect female meiotic progression.

### ATR-mediated signalling controls the synapsis checkpoint

We next assessed the role of ATR in signalling asynapsis. To model the synapsis checkpoint, we used the *Spo11^-/-^*mutant. Meiocytes lacking SPO11 fail to form meiotic DSBs, and as a result exhibit elevated levels of asynapsis, which leads to almost complete loss of primordial oocytes by P21.^10,29^ Depletion of ATR fully rescued the *Spo11^-/-^* oocyte loss phenotype. Oocyte numbers in *Spo11^-/-^ Atr^fl/-^ Stra8-GFPCre* animals were similar to control at P21 (**Figure 2A, B**). While HORMAD2 was retained along asynapsed axes in *Spo11^-/-^ Atr^fl/-^ Stra8-GFPCre* oocytes, the accompanying γH2AX staining typical of *Spo11^-/-^* oocytes was not observed, indicating that asynapsis is still detected but no longer signalled (**Figure 2C**).^30^

**Figure 2.**
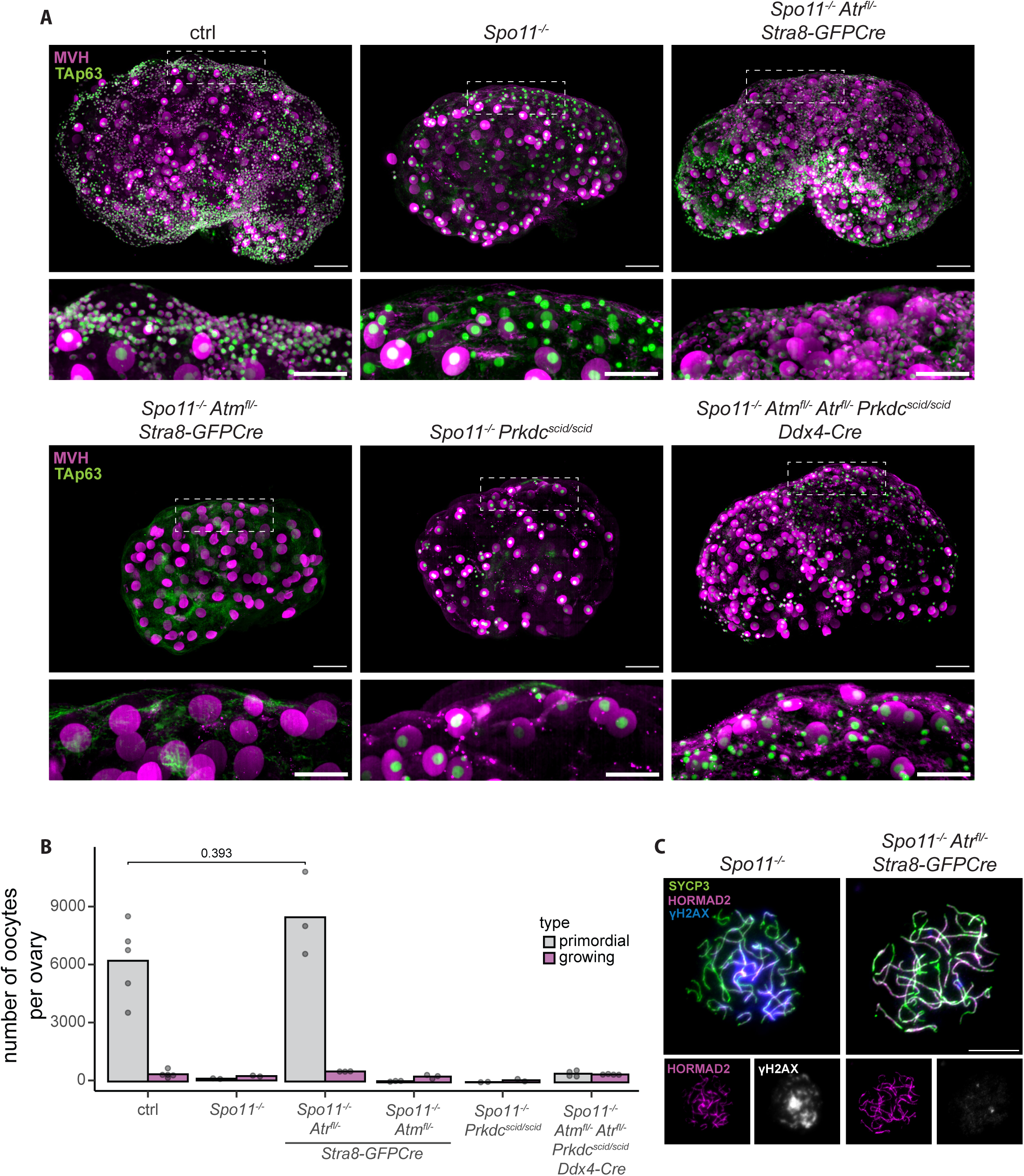
ATR signalling mediates the synapsis checkpoint. **A)** Wholemount ovary images from control, *Spo11^-/-^*, *Spo11^-/-^ Atr^fl/-^ Stra8-GFPCre*, *Spo11^-/-^ Atm^fl/-^ Stra8-GFPCre*, *Spo11^-/-^ Prkdc^scid/scid^* and *Spo11^-/-^ Atm^fl/-^ Atr^fl/-^ Prkdc^scid/scid^ Ddx4-Cre* mice at P21, stained with antibodies against MVH (magenta) and TAp63 (green). Top: z-projection of whole ovaries; scale bar represents 200μm. Bottom: zoomed panels of the boxed areas in top, focusing on the ovarian cortex. Scale bar represents 100μm. **B)** Bar plot showing quantification of oocytes in primordial and growing follicles in all genotypes from (**A**). Bar height represents average number of oocytes. Dots represent counts from individual ovaries. Number of animals used: 5 for control, 2 for *Spo11^-/-^*, 3 for *Spo11^-/-^ Atr^fl/-^ Stra8-GFPCre*, 3 for *Spo11^-/-^ Atm^fl/-^ Stra8-GFPCre*, 2 for *Spo11^-/-^ Prkdc^scid/scid^*and 4 for *Spo11^-/-^ Atm^fl/-^ Atr^fl/-^ Prkdc^scid/scid^ Ddx4-Cre*. p values were calculated using the Mann Whitney U test. Only the comparison between control and rescued *Spo11^-/-^ Atr^fl/-^ Stra8-GFPCre* ovaries is shown. **C)** Chromosome spreads from newborn (P0) *Spo11^-/-^* and *Spo11^-/-^ Atr^fl/-^ Stra8-GFPCre* mice, stained for SYCP3 (green), HORMAD2 (magenta) and γH2AX (blue). Top: Overlay of all channels. Bottom: individual panels showing separate staining of HORMAD2 and γH2AX (here in grey). Scale bar represents 10μm. For *Spo11^-/-^*, 48/51 oocytes (from 1 animal) tested exhibited excessive γH2AX signal; for *Spo11^-/-^ Atr^fl/-^ Stra8-GFPCre*, 11/126 oocytes (from 1 animal) showed limited γH2AX signal.

To verify that ATR is the only PIKK responsible for asynapsis-related signalling, we deleted ATM and DNA-PK in *Spo11^-/-^* females. We observed no rescue in *Spo11^-/-^ Atm^fl/-^ Stra8-GFPCre* or *Spo11^-/-^ Prkdc^scid/scid^* (**Figure 2A, B**), indicating that elimination of *Spo11*-null oocytes depends specifically on ATR.

Next, we sought to define the developmental window during which ATR activates the synapsis checkpoint in *Spo11^-/-^* mutants. Whereas *Stra8-GFPCre*–mediated excision results in ATR protein loss from E15.5 (**Figure S3A**), *Ddx4-Cre*-mediated PIKK depletion occurs later, around P0 (**Figure S1A, B**). In contrast to the complete rescue achieved by early, *Stra8-GFPCre*-mediated ATR deletion, later *Ddx4-Cre*-mediated deletion produced only a minor oocyte rescue, even when combined with loss of both ATM and DNA-PK (*Spo11^-/-^ Atm^fl/-^ Atr^fl/-^ Prkdc^scid/scid^ Ddx4-Cre*) (**Figure 2A, B**). These observations demonstrate that ATR-dependent checkpoint signalling is established before birth, when effective *Ddx4-Cre*-mediated PIKK depletion is first observed.

### ATR controls oocyte elimination in *Dmc1^-/-^* females

We next assessed whether ATR also mediates oocyte elimination in *Dmc1^-/-^*females. *Dmc1*-null oocytes exhibit severe asynapsis and defective recombination, accompanied by extensive DNA damage during late prophase I and complete oocyte loss shortly after birth.^10,31,32^ Early depletion of ATR fully rescued this phenotype, restoring oocyte numbers in *Dmc1^-/-^ Atr^fl/-^ Stra8- GFPCre* ovaries to wild-type levels (**Figure 3A, B**). RPA2 foci persisted in *Dmc1^-/-^ Atr^fl/-^ Stra8-GFPCre* oocytes from P0 ovaries, demonstrating that rescue was not due to early DSB resolution (**Figure 3C**). In contrast, neither ATM depletion nor DNA-PK loss rescued oocyte loss, indicating that elimination of *Dmc1*-null oocytes is dependent on ATR. Furthermore, later PIKK depletion using *Ddx4-Cre* had minimal impact on the *Dmc1*-null phenotype (**Figure 3A, B**). Thus, as in *Spo11*-null oocytes, ATR-dependent checkpoint signalling in *Dmc1*-null oocytes is established during foetal development, most likely during the pachynema stage of prophase I.

**Figure 3.**
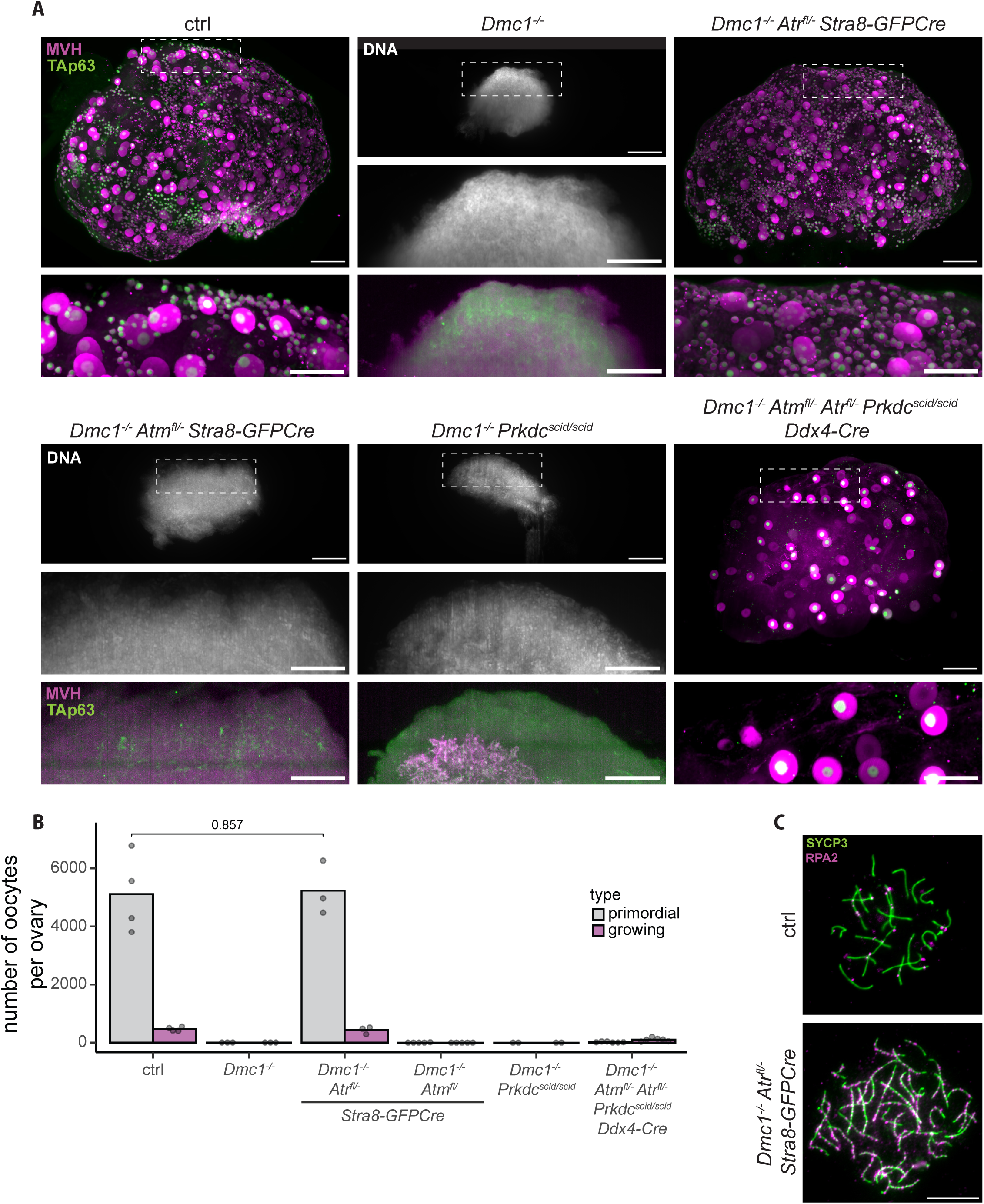
Oocyte death in *Dmc1^-/-^* mutants depends on ATR. **A)** Wholemount ovary images from control, *Dmc1^-/-^*, *Dmc1^-/-^ Atr^fl/-^ Stra8-GFPCre*, *Dmc1^-/-^ Atm^fl/-^ Stra8-GFPCre*, *Dmc1^-/-^ Prkdc^scid/scid^* and *Dmc1^-/-^ Atm^fl/-^ Atr^fl/-^ Prkdc^scid/scid^ Ddx4-Cre* mice at P21, stained with antibodies against MVH (magenta) and TAp63 (green). Top: z-projection of whole ovaries; scale bar represents 200μm. Bottom: zoomed panels of the boxed areas in top, focusing on the ovarian cortex; scale bar represents 100μm. For *Dmc1^-/-^*, *Dmc1^-/-^ Atm^fl/-^ Stra8-GFPCre* and *Dmc1^-/-^ Prkdc^scid/scid^*, DNA (counterstained with propidium iodide; grey) is shown at the top, due to the lack of signal in other channels. **B)** Bar plot showing quantification of oocytes in primordial and growing follicles in all genotypes from (**A**). Bar height represents average number of oocytes. Dots represent counts from individual ovaries. Number of animals used: 4 for control, 3 for *Dmc1^-/-^*, 3 for *Dmc1^-/-^ Atr^fl/-^ Stra8-GFPCre*, 4 for *Dmc1^-/-^ Atm^fl/-^ Stra8-GFPCre*, 2 for *Dmc1^-/-^ Prkdc^scid/scid^* and 6 for *Dmc1^-/-^ Atm^fl/-^ Atr^fl/-^ Prkdc^scid/scid^ Ddx4-Cre*. p values were calculated using the Mann Whitney U test. Only the comparison between control and rescued *Dmc1^-/-^ Atr^fl/-^ Stra8-GFPCre* ovaries is shown. **C)** Chromosome spreads of pachytene and zygotene-like oocytes from newborn (P0) control and *Dmc1^-/-^ Atr^fl/-^ Stra8-GFPCre* mice, stained for SYCP3 (green) and RPA2 (magenta). Scale bar represents 10μm. For control, the median RPA2 foci per oocyte was 18 (as seen in **Figure S3C**); for *Dmc1^-/-^ Atr^fl/-^ Stra8-GFPCre*, the media RPA2 foci per oocyte was 213 (32 oocytes from 1 animal).

### ATR controls oocyte elimination in response to persistent meiotic recombination lesions

Since *Dmc1^-/-^* oocytes exhibit both impaired meiotic recombination and synapsis, it remained unclear which of these two defects triggered ATR-dependent elimination. We therefore sought to generate a model in which meiotic recombination is perturbed while homolog synapsis is unaffected.

Conditional depletion of ATM offered a potential means of achieving this separation. ATM has important early functions in meiotic DSB homeostasis, and constitutive *Atm^-/-^*oocytes exhibit substantially elevated DSB levels and defective synapsis.^24,33,34^ We reasoned that delaying ATM depletion until after meiotic entry might circumvent these early defects while revealing later recombination phenotypes. We therefore depleted ATM using *Stra8-GFPCre*. ATM protein levels were comparable between control and mutant ovaries at E13.5, but reduced in mutant ovaries by E15.5 (**Figure S4A**), confirming effective depletion during meiotic prophase.

We examined DSB formation and repair in the resulting conditional ATM-deficient oocytes. At E16.5, *Atm^fl/-^ Stra8-GFPCre* pachytene oocytes contained RPA2 focus counts comparable to controls, indicating that early DSB homeostasis was largely preserved. However, by P0, mutant pachytene oocytes exhibited elevated numbers of RPA2 foci, consistent with delayed resolution of meiotic recombination intermediates (**Figure 4A-D**). Despite this persistence of recombination-associated lesions, the frequency of HORMAD2-positive pachytene oocytes was comparable between mutant and control ovaries, indicating largely normal homologue synapsis (**Figure S4B, C**). Conditional ATM depletion therefore provides a model in which persistent meiotic recombination intermediates occur in the absence of increased asynapsis.

**Figure 4.**
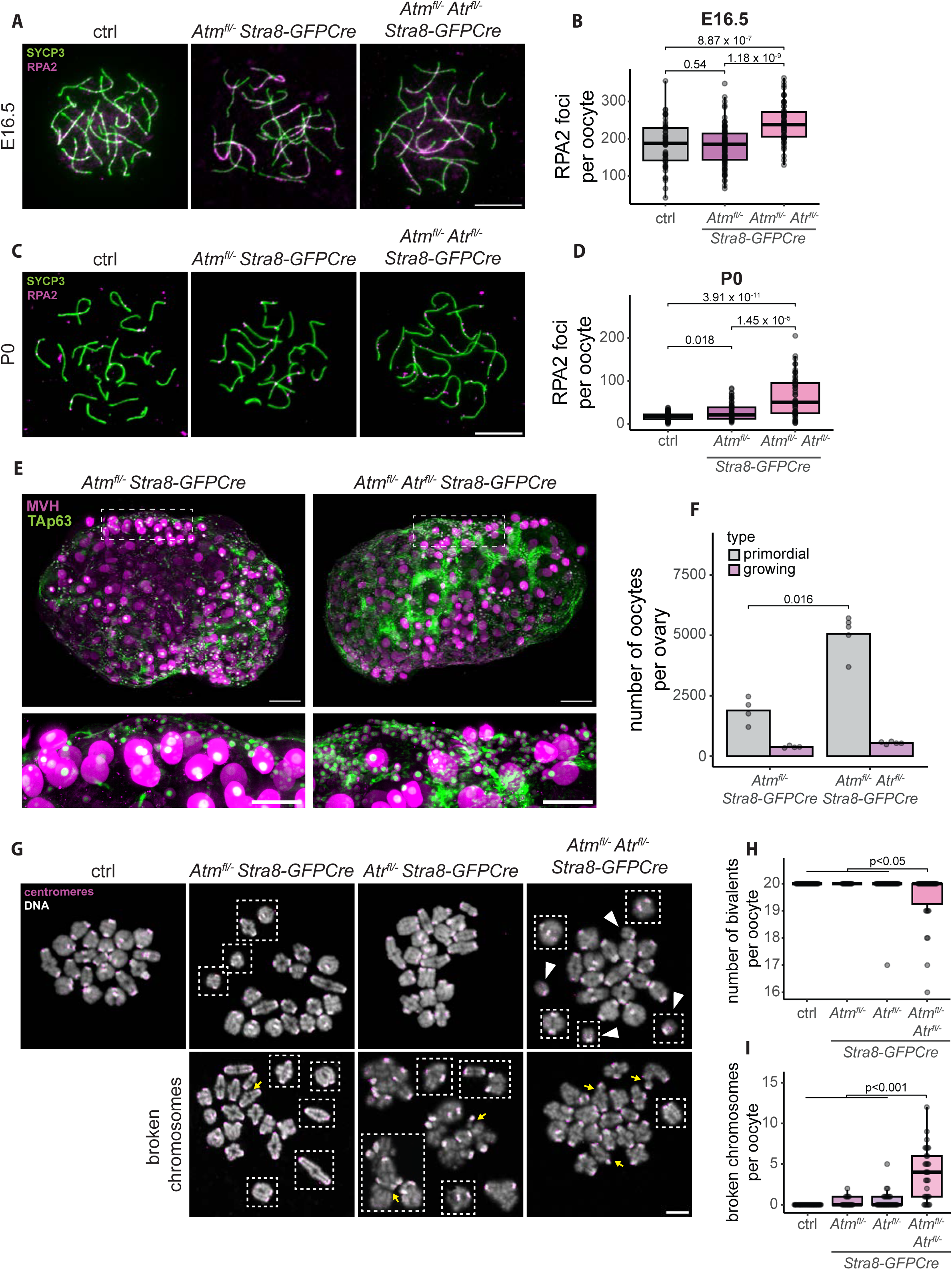
ATR signalling mediates the recombination checkpoint. **A)** Chromosome spreads from control, *Atm^fl/-^ Stra8-GFPCre* and *Atm^fl/-^ Atr^fl/-^ Stra8-GFPCre* E16.5 embryos, stained for SYCP3 (green) and RPA2 (magenta). **B)** Box and whiskers plot of quantified RPA2 foci in genotypes from (**A**). Dots represent individual cells. 53 (from 3 females), 88 (from 3 females) and 90 (from 3 females) oocytes were assessed, respectively. **C)** Chromosome spreads from control, *Atm^fl/-^ Stra8-GFPCre* and *Atm^fl/-^ Atr^fl/-^ Stra8-GFPCre* P0 females, stained for SYCP3 (green) and RPA2 (magenta). **D)** Box and whiskers plot of quantified RPA2 foci in genotypes from (**C**). Dots represent individual cells. 72 (from 3 females), 66 (from 3 females) and 56 (from 3 females) oocytes were assessed, respectively. Scale bars in (**A**) and (**C**) indicate 10μm. p values in (**B**) and (**D**) were calculated using the Kruskal-Wallis test, followed by Dunn’s post hoc test with Benjamini-Hochberg correction. **E**) Wholemount images of P21 ovaries from *Atm^fl/-^ Stra8-GFPCre* and *Atm^fl/-^ Atr^fl/-^ Stra8-GFPCre* females, stained for MVH (magenta) and TAp63 (green). Top: z-projections of whole ovaries. Scale bar represents 200μm. Bottom: zoomed panels of the boxed areas from top, focusing on the cortex of the ovary. Scale bar indicates 100μm. **F)** Bar plot showing the average count of primordial and growing follicles of genotypes from (**E**). Dots represent counts from individual animals. Number of females used: 4 for *Atm^fl/-^ Stra8-GFPCre* and 5 for *Atm^fl/-^ Atr^fl/-^ Stra8-GFPCre*. p value was calculated using the Mann Whitney U test. **G)** Chromosome spreads from metaphase I oocytes of control, *Atm^fl/-^ Stra8-GFPCre*, *Atr^fl/-^ Stra8-GFPCre* and *Atm^fl/-^ Atr^fl/-^ Stra8-GFPCre* females, stained with CREST serum (magenta) and counterstained with Hoechst 33342 (DNA, grey). Dashed white boxes indicate chromosomes that were outside of the field of view, which were separately pasted on empty space. Top row shows oocytes without DNA damage, and for *Atm^fl/-^ Atr^fl/-^ Stra8-GFPCre* an example of achiasmate chromosomes, indicated by white arrowheads. Bottom row shows examples of oocytes with broken chromosomes, indicated by yellow arrows. Scale bar represents 10μm. **H)** Box and whiskers plot showing the number of homologues that have formed chiasmata and appear as bivalents in genotypes from (**G**). Dots represent individual oocytes. 26 (from 3 females), 12 (from 3 females), 27 (from 3 females), and 30 (from 3 females) oocytes were assessed, respectively. **I)** Box and whiskers plot showing the number of broken chromosomes per oocyte in the same pool of oocytes as (**H**). Dots represent individual oocytes. p values in (**H**) and (**I**) were calculated using the Kruskal-Wallis test, followed by Dunn’s post hoc test with Benjamini-Hochberg correction.

Next, we asked whether these persistent recombination-associated lesions trigger ATR-dependent oocyte elimination. At P21, *Atm^fl/-^ Stra8-GFPCre* ovaries contained fewer primordial oocytes than controls (**Figure 4E, F** and **Figure S3F, G**). Importantly, co-depletion of ATR fully restored oocyte numbers. Thus, elimination of *Atm^fl/-^ Stra8-GFPCre* oocytes is ATR-dependent.

We then examined meiotic progression in the rescued *Atm^fl/-^ Atr^fl/-^ Stra8-GFPCre* females. Oocytes depleted of both ATM and ATR showed elevated levels of RPA2 foci in pachytene cells from E16.5 ovaries (**Figure 4A, B**), indicating slower DSB processing kinetics. Additionally, pachytene oocytes from P0 ovaries also contained elevated numbers of RPA2 foci compared with both control and ATM-depleted oocytes (**Figure 4C, D**). This could reflect an additional requirement for ATR in the processing or resolution of meiotic DSBs in the absence of ATM, or alternatively the survival of oocytes with DNA damage that would otherwise have been eliminated by ATR-dependent surveillance. Co-depletion of ATM and ATR also increased the frequency of HORMAD2-positive pachytene oocytes, despite synapsis being largely unaffected following depletion of either kinase alone (**Figure S4B, C**). This suggests that ATM and ATR have partially redundant functions in supporting efficient homologue synapsis.

At metaphase I, 8/30 (26.7%) rescued *Atm^fl/-^ Atr^fl/-^ Stra8-GFPCre* oocytes contained one or more achiasmate chromosomes, indicating defective crossover formation (**Figure 4G, H**). In addition, visible chromosome breaks were present in 83.3% of rescued oocytes (**Figure 4G, I**). Thus, ATR depletion revealed the later consequences of defective meiotic recombination, allowing oocytes with failed crossover formation and persistent chromosome damage to survive to metaphase I. Together, these findings demonstrate that ATR controls oocyte elimination in response to persistent recombination-associated DNA lesions independently of homolog asynapsis.

## Discussion

Here, we establish the roles of the individual PIKK kinases ATM, ATR and DNA-PK in the mammalian oocyte checkpoints (**Figure 6**). We show that ATR is the critical PIKK driving both the synapsis and recombination checkpoints. This contrasts with the canonical somatic DNA damage response, in which DSB signalling is primarily mediated by ATM.^35^ The preferential use of ATR during meiotic prophase may reflect the unusual substrates that must be monitored during recombination: programmed DSBs are extensively resected to generate RPA-coated ssDNA, providing a potent platform for ATR recruitment and activation.^36–39^ Subsequently, in postnatal oocytes, DNA damage checkpoint signalling switches to an ATM-dominated response. Thus, oogenesis involves a developmental reconfiguration of DNA damage signalling, from ATR-dependent surveillance of meiotic synapsis and recombination to canonical ATM-dependent surveillance of DNA damage.

**Figure 5.**
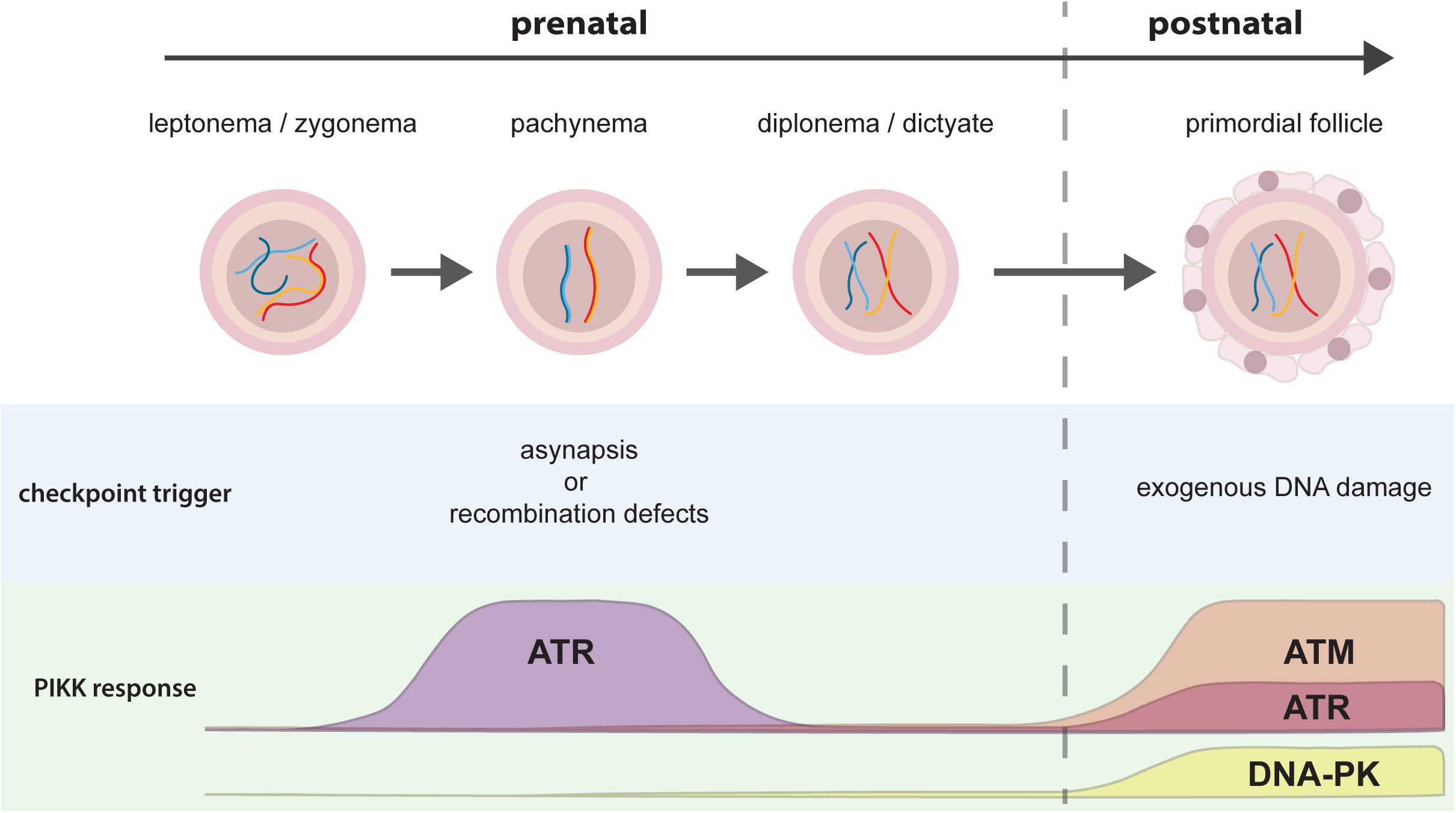
Schematic representation of PIKK involvement in prenatal and postnatal oocyte checkpoints.

ATR is required for both the synapsis and recombination checkpoints, which might imply that these two defects converge on a common downstream elimination pathway. However, previous work has shown that they engage genetically separable downstream apoptotic programmes: loss of PUMA/NOXA or BAX rescues oocyte elimination caused by defective recombination, but not that caused by synapsis failure in *Spo11^−/−^* oocytes.^40^ This is further supported by the differential timing of oocyte elimination between *Dmc1^-/-^* and *Spo11^-/-^*females.^10^ How ATR distinguishes between these different meiotic defects and channels them into distinct downstream effector programmes is an interesting area for future study.

Although ATR is the principal checkpoint kinase in foetal oocytes, ATM is also active during meiotic prophase, where it regulates DSB formation and processing.^24,33,34^ An important unresolved question is how such ATM activity does not trigger the oocyte elimination pathway. In somatic cells, ATM can activate CHK2, leading ultimately to phosphorylation and stabilisation of p53.^41–44^ At what level this signalling axis is restrained during meiotic prophase remains unclear. Regulation could occur upstream, by limiting transmission of the ATM signal to CHK2, downstream through suppression of p53, or at multiple points in the pathway. Interestingly, p53 is not detectable during early stages of ovarian development.^45^ Clarifying the mechanisms that selectively control ATM-mediated responses to DSBs could inform potential strategies for selective protection of healthy cells during genotoxic treatments.

## Resource availability

The code for the oocyte segmentation analysis has been deposited and is publicly available at Zenodo: https://doi.org/10.5281/zenodo.23249701

### Lead contact

Further information and resource requests should be directed to and will be fulfilled by the lead contact, James Turner.

### AI usage

During the preparation of this work, the authors used AI tools (Claude, ChatGPT) for grammar and syntax checks, occasional re-phrasing of convoluted sentences, and for a prototype of the schematic in Figure 6, following the authors’ design. All content was reviewed, edited and verified by the authors, who take full responsibility for it.

## Acknowledgements

We would like to thank the Biological Research Facility of the Francis Crick Institute for their help in maintaining our mouse colonies, the Experimental Histopathology facility for their contributions and the Advanced Light Microscopy facility for their expertise and support. We thank the members of the J.M.A.T. lab for their feedback and discussions on the manuscript. This work was supported by core funding of the Francis Crick Institute, which is received from Cancer Research UK (CC2052), the UK Medical Research Council (CC2052), and the Wellcome Trust (CC2052).

## Author contributions

J.M.A.T. conceived the project. G.S. and E.E. performed the irradiation experiments, immunostaining of ovary sections and oocyte quantifications. G.S. and A.P.Z. performed GV-oocyte harvests, calculated polar body extrusion rates and performed metaphase II oocyte spreads. G.S. and M.J.P. performed oocyte prophase spreads and quantified RPA2 foci. G.S. performed ovary harvests, wholemount immunostaining and clearing of ovaries, oocyte segmentation and quantification; immunostaining and fragment quantification of metaphase II spreads; oocyte harvests, preparation of metaphase I spreads, immunostaining and quantification of broken chromosomes; HORMAD2 immunostainings and quantification; western blot analyses; zygote harvests, *in vitro* culture and quantification of embryo development. T.F. and V.D. developed the pipeline for oocyte segmentation from wholemount images. O.O. maintained mouse colonies and generated mutant combinations. G.S. and J.M.A.T. wrote the manuscript.

---

**Figure S1.**
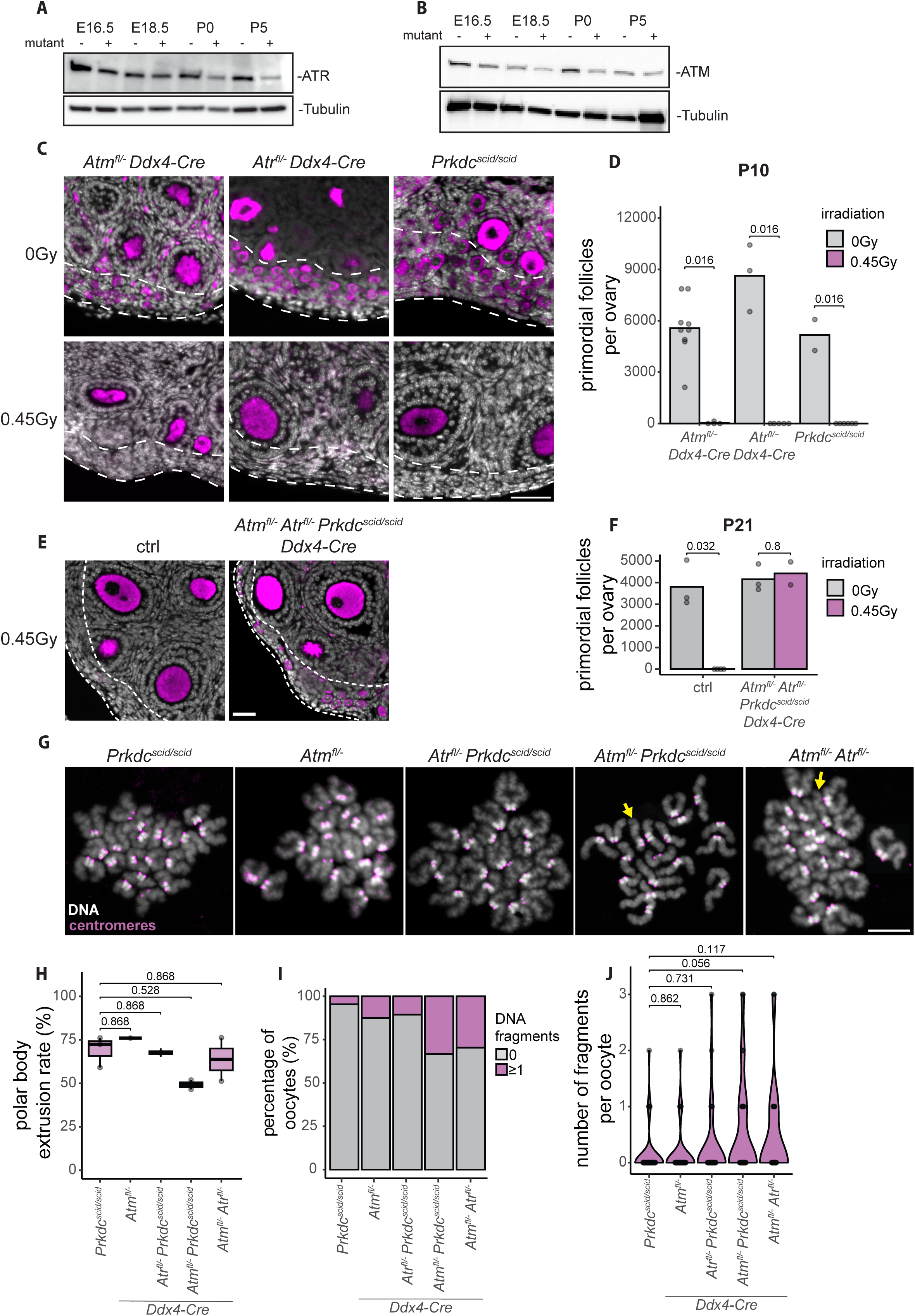
Roles of ATM, ATR and DNA-PK in postnatal DNA damage response and DSB repair. **A, B)** Western blot analysis of protein levels of ATR and ATM in ovaries from control (marked as mutant “-”) and *Atm^fl/-^ Atr^fl/-^ Prkdc^scid/scid^ Ddx4-Cre* mice (marked as mutant “+”) at E16.5, E18.5, P0 and P5, respectively. 4-6 ovaries were used per lane. Tubulin was used as an internal control. **C)** Widefield images of ovary sections from *Atm^fl/-^ Ddx4-Cre*, *Atr^fl/-^ Ddx4- Cre* and *Prkdc^scid/scid^* P10 females without (0Gy) and after 0.45Gy of irradiation treatment at P5, stained with MVH (magenta) and counterstained with DAPI (DNA, grey). Scale bar represents 50μm. **D)** Bar plot representing the average number of primordial follicles of genotypes and conditions shown in (**c**). Dots represent oocyte counts from individual animals. Number of females used: 9 (0Gy) and 4 (0.45Gy) for *Atm^fl/-^ Ddx4-Cre*, 3 (0Gy) and 5 (0.45Gy) for *Atr^fl/-^ Ddx4-Cre*, 2 (0Gy) and 6 (0.45Gy) for *Prkdc^scid/scid^* **E)** Widefield images of sections of P21 ovaries from control and *Atm^fl/-^ Atr^fl/-^ Prkdc^scid/scid^ Ddx4-Cre* females after 0.45Gy of irradiation treatment at P5, stained with MVH (magenta) and counterstained with DAPI (DNA, grey). Dashed white lines in (**C**) and (**E**) indicate the ovarian cortex. Scale bar represents 50μm. **F)** Bar plot of average number of primordial follicles from genotypes shown in (**E**). Dots represent counts from individual animals. Number of females used: 3 (0Gy) and 4 (0.45Gy) for control, 3 (0Gy) and 2 (0.45Gy) for *Atm^fl/-^ Atr^fl/-^ Prkdc^scid/scid^ Ddx4-Cre*. p values were calculated using the Mann Whitney U test. **G)** Chromosome spreads of metaphase II oocytes from *Prkdc^scid/scid^*, *Atr^fl/-^ Prkdc^scid/scid^ Ddx4-Cre*, *Atm^fl/-^ Prkdc^scid/scid^ Ddx4-Cre*, *Atm^fl/-^ Atr^fl/-^ Ddx4- Cre* and *Atm^fl/-^ Ddx4-Cre*, stained using CREST serum (magenta) and counterstained with Hoechst 33342 (DNA, grey). Yellow arrows point to chromosome fragments. **H)** Box and whiskers plot of polar body extrusion rates for oocytes from genotypes shown in (**G**). 97 (from 3 females), 105 (from 2 females), 70 (from 2 females), 121 (from 2 females) and 50 (from 1 female) oocytes were assessed. **I**) Stacked bar plot of percentage of oocytes that have no, or at least one chromosome fragment, from genotypes shown in (**G**). **J)** Violin plot showing the number of fragments in the same pool of oocytes as (**I**). 44 (from 5 females), 38 (from 3 females), 36 (from 3 females), 27 (from 3 females) and 48 (from 4 females) oocytes were assessed. p values in (**H**) and (**J**) were calculated using the Kruskal-Wallis test, followed by Dunn’s post hoc test with Benjamini-Hochberg correction.

**Figure S2.**
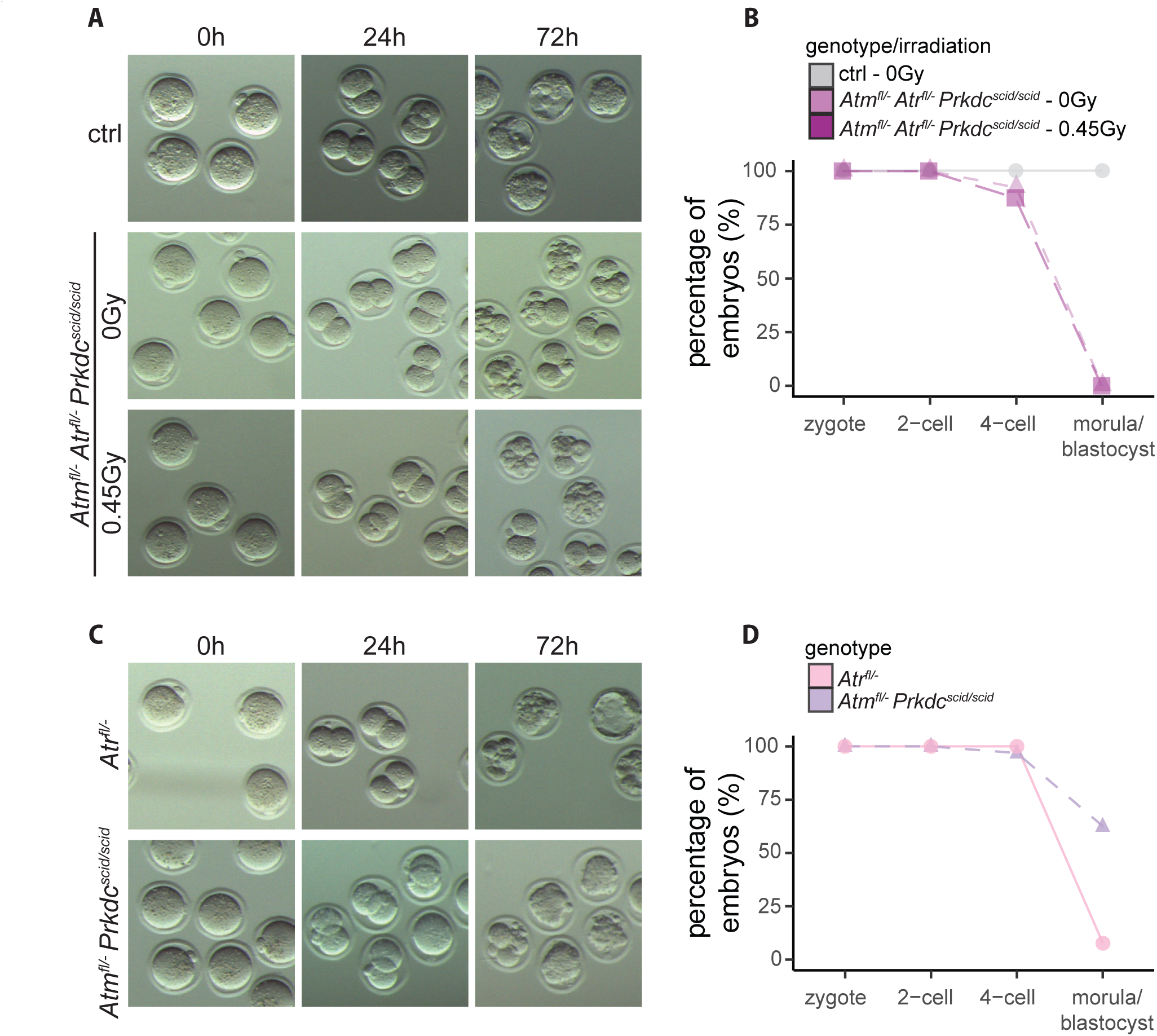
Fertility of different PIKK mutants. **A)** Images of cultured embryos from control and *Atm^fl/-^ Atr^fl/-^ Prkdc^scid/scid^ Ddx4-Cre* (non-irradiated and irradiated at P5) mothers at day of harvest (0h), and after 24 and 72 hours of culture. **B)** Line graph summarizing the percentage of zygotes that achieved the 2-cell, 4-cell and morula or blastocyst stage, from each genotype shown in (**A**). Percentages were calculated from 66 (from 7 mothers), 69 (from 7 mothers) and 46 (from 5 mothers) embryos, respectively. **C)** Images of cultured embryos from *Atr^fl/-^ Ddx4- Cre* and *Atm^fl/-^ Prkdc^scid/scid^ Ddx4-Cre* mothers at day of harvest (0h), as well as after 24 and 72 hours of culture. **D)** Line graph summarising the percentage of zygotes that achieved the 2-cell, 4-cell and morula or blastocyst stage, from each genotype shown in (**D**). 34 (from 4 mothers) and 23 (from 3 mothers) embryos were used for percentage calculations, respectively.

**Figure S3.**
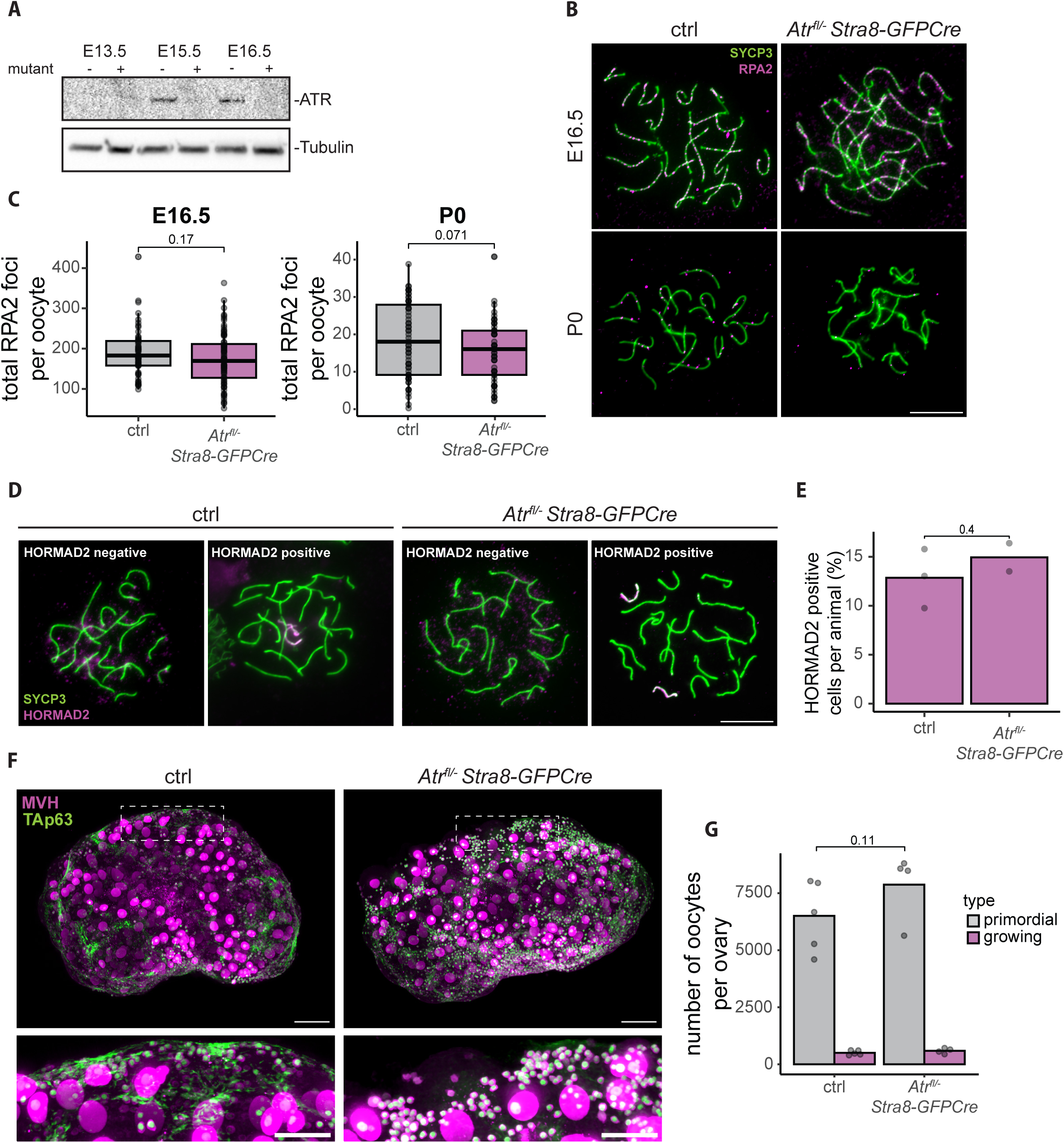
ATR roles in female meiosis. **A)** Western blot analysis of ATR protein levels in control (marked as mutant “-”) and *Atr^fl/-^ Stra8-GFPCre* (marked as mutant “+”) ovaries from E13.5, E15.5 and E16.5 embryos. Between 4 and 6 ovaries were used for each timepoint; Tubulin was used as an internal control. **B)** Oocyte chromosome spreads from E16.5 and P0 control and *Atr^fl/-^ Stra8-GFPCre* ovaries, stained for SYCP3 (green) and RPA2 (magenta). Scale bar indicates 10μm. **C)** Box and whiskers plot of quantification of RPA2 foci from control and *Atr^fl/-^ Stra8-GFPCre* oocytes shown in (**B**), at E16.5 and P0. E16.5: number of oocytes assessed was 57 (from 3 females) and 76 (from 3 females), respectively. P0: number of oocytes was 61 (from 4 females) and 59 (from 4 females) oocytes, respectively. **D)** Oocyte chromosome spreads from E18.5 control and *Atr^fl/-^ Stra8-GFPCre* ovaries, stained for SYCP3 (green) and HORMAD2 (magenta). Examples of HORMAD2 negative and positive oocytes are shown for both genotypes. Scale bar indicates 10μm. **E)** Bar plot showing the percentage of HORMAD2-positive pachytene cells in control and *Atr^fl/-^ Stra8-GFPCre* oocytes. Each dot represents the percentage of positive cells from one animal. 83 (from 3 females) and 135 (from 2 females) cells were analysed, respectively. **F)** Wholemount immunofluorescent images of control and *Atr^fl/-^ Stra8-GFPCre* ovaries harvested at P21. Top: whole ovaries stained for MVH (magenta) and TAp63 (green). Scale bars indicate 200μm. Bottom: zoomed panels on the boxed areas in top, focusing on the ovarian cortex. Scale bars represent 100μm. **G)** Bar plots of quantified numbers of oocytes from primordial and growing follicles in genotypes shown in (**F**). Bar height represents the average number of oocytes from multiple animals. Dots represent counts from individual females. Number of females used: 5 for control, 4 for *Atr^fl/-^ Stra8-GFPCre*. p values in (**C**), (**E**) and (**G**) were calculated using the Mann-Whitney U test.

**Figure S4.**
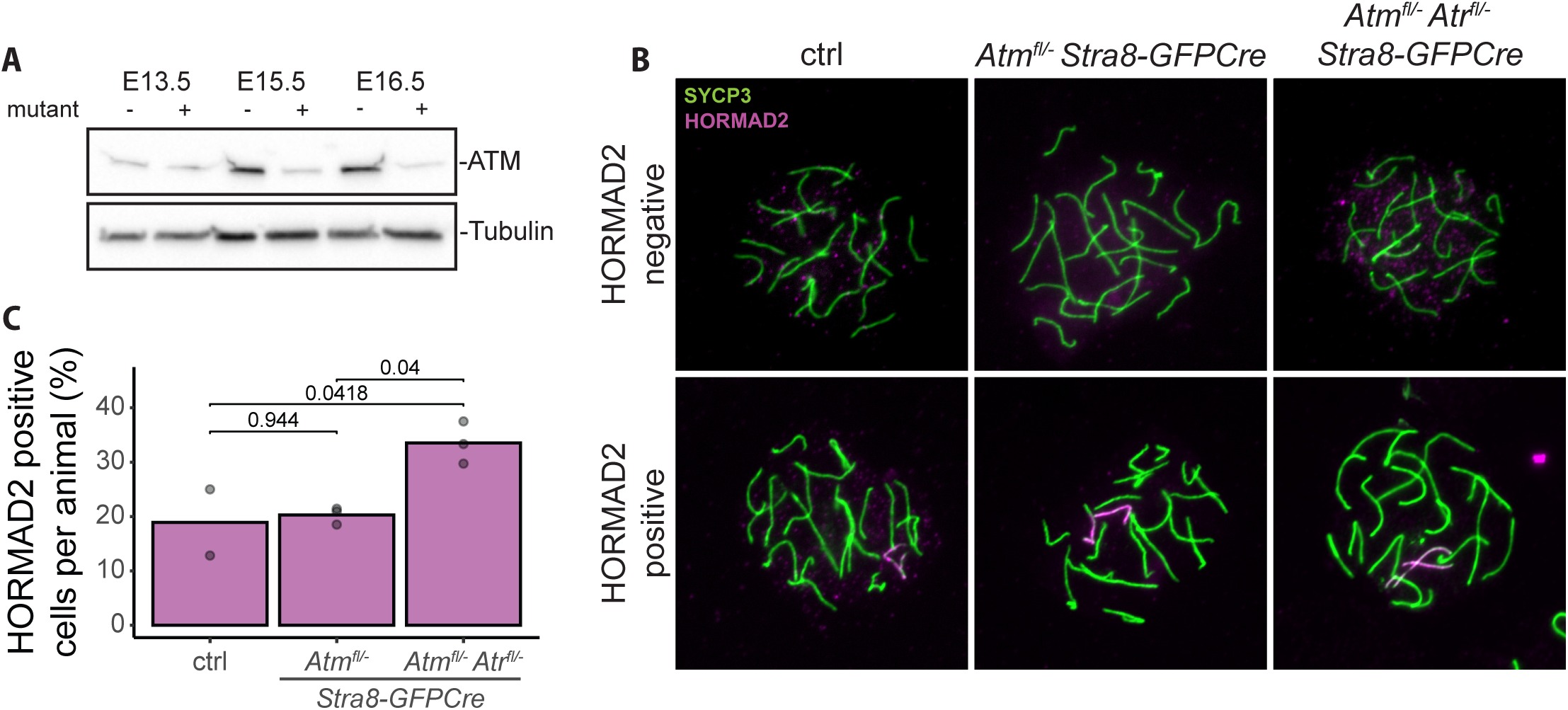
Asynapsis levels in *Atm^fl/-^ Stra8GFP-Cre* and *Atm^fl/-^ Atr^fl/-^ Stra8GFP-Cre* oocytes. **A)** Western blot analysis of ATM protein levels in control (marked as mutant “-”) and *Atm^fl/-^ Stra8-GFPCre* ovaries (marked as mutant “+”) from E13.5, E15.5 and E16.5 embryos. Between 4 and 6 ovaries were used per lane; Tubulin was used as a loading control. **B)** Oocyte chromosome spreads from control, *Atm^fl/-^ Stra8GFP-Cre* and *Atm^fl/-^ Atr^fl/-^ Stra8GFP-Cre* ovaries, harvested at E18.5 and stained for SYCP3 (green) and HORMAD2 (magenta). Examples of HORMAD2-negative and -positive cells are shown for all genotypes. Scale bar indicates 10μm. **C)** Bar plot of quantified percentages of pachytene oocytes with detectable HORMAD2 signal in genotypes shown in (**B**). 59 (from 2 females), 84 (from 3 females) and 76 (from 3 females) cells were assessed, respectively. p values were calculated using the Kruskal-Wallis test, followed by Dunn’s post hoc test with Benjamini-Hochberg correction.

## Methods

### Mouse colony maintenance and irradiation of pups

Mouse lines (*Mus musculus*, C57BL/6J strain) were maintained under UK Home Office Regulations, UK Animals (Scientific Procedures) Act 1986, and in line with ethical guidelines of the Francis Crick Institute. Permission for animal experiments was granted by The Crick Biological Research Facility Strategic Oversight Committee (BRF-SOC) incorporating Animal Welfare and Ethical Review Body (AWERB) (project licence PP1234304). Mice were housed in individually ventilated cages with free access to food and water, at a constant temperature of 20-22°C and humidity of 49-62%. The mice were kept at a 12h-light/12h-dark cycle. Mouse lines were checked daily. Genetically modified models used were previously published and were predominantly kept on a C57BL/6J background: *Atm*,^33^ *Atm flox*,^48^ *Atr*,^49^ *Atr flox*,^50^ *Ddx4-Cre*,^25^ *Stra8-GFPCre*,^51^ *Spo11*,^29^ *Dmc1*,^31^ *Prkdc scid*.^52^ For irradiation experiments, P5 pups were exposed to a single low dose (0.45Gy) of ionising radiation using a GSR-D1 ^137^Caesium irradiator.

### Ovary sectioning and immunofluorescence

Ovaries collected from P10 and P21 mice were fixed in 4% paraformaldehyde (PFA, diluted in PBS) overnight, shaking at 4°C, followed by PBS washes, paraffin embedding and serial sectioning at 6μm. For immunofluorescence, slides were deparaffinised and rehydrated using xylene and serial ethanol washes, followed by antigen retrieval by boiling for 30 minutes in 0.1M sodium citrate buffer. Blocking was performed for 15 minutes in 5% bovine serum diluted in PBS supplemented with 0.1% Tween 20 (PBS-T) at room temperature (RT). Primary antibody (rabbit anti-MVH/Ddx4, ab13840 Abcam, 1:100 dilution) was diluted in blocking buffer and was incubated on slides overnight at 37°C. Secondary antibody (Alexa Fluor®488, 1:200) was diluted in PBS-T, which also included 4’,6-diamidino-2-phenylindole (DAPI). Secondary incubation was performed at room temperature for 1 hour. Slides were mounted with VECTASHIELD Antifade Mounting Medium (H-1000, Vector Laboratories). Imaging was performed using a Zeiss Observer.Z1 inverted microscope with a 20X objective lens, or an Olympus VS120 Slide Scanner with a U-HGLGPS mercury lamp and an Olympus UPlanApo 20x/1.35 NA objective. Images were captured through a 2/3” CCD camera and analysed using ImageJ.

### Wholemount ovary preparation, immunofluorescence and clearing

For experiments assessing meiotic checkpoint control (referring to Figures 2-4 and their supplements), a wholemount imaging approach was adopted. Ovaries harvested from P21 animals were fixed in 4% PFA overnight at 4°C, washed with PBS and dehydrated using serial dilutions of methanol in H_2_O, before being stored in 100% methanol at -20°C.

The iDISCO protocol was used for immunofluorescence and clearing, as described for ovaries previously.^53^ In short, dehydrated, stored ovaries were pre-treated with a 66% dichloromethane (270997, Sigma-Aldrich)/33% methanol solution overnight at RT, before being bleached with 5% H_2_O_2_ solution in methanol overnight at 4°C. Following this, ovaries were rehydrated using serial dilutions of methanol/H_2_O, permeabilised and blocked. Primary antibodies used were always rabbit anti-MVH (ab13840, Abcam, 1:100 dilution) and mouse anti-TAp63 (NBP3-11703, Novus Biologicals, 1:250 dilution). Primary incubation was performed for 4 days at 37°C without shaking. Ovaries were then incubated with secondary antibodies (anti-mouse Alexa Fluor® 647 and anti-rabbit Alexa Fluor®790, at 1:500 and 1:100 dilutions, respectively) for 3 days at 37°C. In the secondary antibody incubation, propidium iodide at 1μg/ml was also included to counterstain nucleic acids. Ovaries were then embedded in blocks of 1% low-melting agarose, dehydrated using serial methanol/H_2_O dilutions, washed with a 66% dichloromethane/33% methanol solution for 3 hours and two final washes with 100% dichloromethane. For clearing, agarose blocks were moved into di-benzyl ether (108014, Sigma-Aldrich). All steps performed at 37°C used PBS supplemented with 0.01% Sodium Azide (S2002, Sigma-Aldrich).

Imaging was performed using a LaVision BioTec UltraMiscroscope II light-sheet microscope (Miltenyi Biotec), equipped with an Olympus MVPLAPO 2X/0.5 objective and a detachable LaVision BioTech dipping cap, at a total magnification of 8X. Image acquisition was done using an Andor Zyla 5.5 camera (Oxford Instruments). Whole ovaries were scanned every 2μm using 50% laser width and 3.4μm laser thickness.

### Wholemount ovary oocyte segmentation and quantification

Three-dimensional segmentation was performed in Fiji, using Cellpose and Trackmate plugins. The MVH signal was used for segmentation of objects of two different pixel sizes, 15 and 60 pixels. These sizes were picked because their sum resulted in the best segmentation of both primordial and growing follicles of variable sizes. 2D Cellpose segmentations were merged into a 3D segmentation by converting the Z plane (number of slices in Z) to time. This allowed the use of Trackmate to construct the 3D objects.^54^ Segmentation results from Trackmate were manually adjusted for each sample, using size, circularity and contrast scores. Results were exported as label images, the two segmentation maps of each ovary were merged and overlapping segmentations were removed using a custom python script developed by the Crick Advanced Light Microscopy team. Object statistics were obtained by importing label images in Imaris (v9.0, Oxford Instruments) 3D reconstructions of ovaries. Further filtering was performed using area, volume and sphericity scores to segment oocytes and categorise them in “primordial” and “growing” populations.

### Oocyte prophase spread preparation, immunofluorescence and imaging

Oocyte prophase I spreads were performed as previously published.^11^ In short, ovaries from E16.5 and E18.5 embryos, and P0 pups, were harvested and cleaned of surrounding tissues in PBS, before being submerged in 20μl of hypotonic extraction buffer (30 mM Tris–HCl, 17 mM trisodium citrate dihydrate, 5 mM EDTA, 50 mM sucrose, 0.5 mM DTT, 0.5 mM PMSF, supplemented with 1× Protease Inhibitor Cocktail). E16.5 ovaries were incubated for 5 minutes, E18.5 for 10 minutes and P0 for 15 minutes, all at RT. After incubation, the buffer was removed and 15μl of 100mM sucrose in 5mM sodium borate solution (J62902, ThermoFischer) were added. Ovaries were punctured to release the oocytes using 30G needles. Big chunks of tissue were removed and 9μl of 65mM sucrose in 5mM sodium borate solution were added, mixed by pipetting and incubated for 3 minutes. After the last incubation, 1.5μl of the cell suspension were moved into wells drawn by lipophilic marker on SuperFrost Plus^TM^ Adhesion Microscope slides (enhanced version, J1840AMNZ, Epredia), containing 20μL of fixative solution (1% PFA, 50mM sodium borate buffer pH 9.2, 0.15% Triton X-100). Cells were fixed for 45 minutes in a wet chamber, and slides were air dried in a fume hood. Once fully dried, slides were washed in 0.4% Kodak Photo-Flo (1464510, Kodak) for 5 minutes, followed by 5 minutes in H_2_O and further air-drying. Dried slides were stored at - 80°C.

For immunofluorescence, slides were thawed for 5 minutes in PBS, followed by 5 minutes in PBS supplemented with 0.05% Triton X-100 (PBS-Tx). Slides were subsequently incubated with blocking buffer (2.5% BSA, 0.05% Tween 20 in PBS) for 1 hour at RT. Primary antibodies were diluted in blocking solution and incubation was done overnight at 37°C. Primary antibodies used: anti-SYCP3 (guinea pig host, produced in house, dilution 1:500), anti-RPA32/2 (rabbit host, Abcam ab10359, dilution 1:100), anti-HORMAD2 (rabbit host, a gift from Attila Tóth, dilution 1:3000) and anti-γH2AX (mouse host, Millipore 05-636, dilution 1:1000). After incubation with primary antibodies, slides were washed 3 x 5 minutes with PBS-Tx. Secondary antibodies were diluted in blocking solution and were incubated with slides for 1.5 hours at RT. Secondaries used: anti-guinea pig Alexa Fluor 488, anti-rabbit Alexa Fluor 555, anti-mouse Alexa Fluor 647 (ThermoFischer, all diluted 1:500). After 3 x 10-minute washes, slides were mounted in VECTASHIELD Antifade Mounting Medium with DAPI (H-1200, Vector Laboratories).

Imaging was performed using a Zeiss Observer.Z1 inverted microscope with a 63X objective lens.

### Oocyte harvesting, *in vitro* culture, preparation of metaphase spreads, immunofluorescence and imaging

Mature GV-oocytes were harvested from mice aged 5-8 weeks. Ovaries were dissected and cleaned of surrounding tissues in PBS. Then, ovaries were moved into a drop of M2 medium and were punctured using 30G needles to release oocytes. Collection and washes of GV-oocytes were performed by mouth-pipetting. GV-oocytes were washed through 9μl drops of M2 (M7167, Sigma-Aldrich) under tissue culture oil (ART-4008, CooperSurgical) and were cultured at 37°C in M2 to allow for meiosis resumption. For metaphase I spreads, oocytes were cultured for 5.5-7 hours, and for metaphase II for 16-17 hours (overnight). After culture, oocytes were stripped of the zona-pellucida by a short incubation in drops of pre-warmed Tyrode’s acidic solution (T1788, Sigma-Aldrich). Afterwards, oocytes were swollen by incubation in a pre-warmed 1:1 solution of foetal bovine serum (FBS) in H_2_O, for 13 minutes at 37°C, on 0.5% agarose plates, to prevent oocytes from attaching to the bottom of the dish. Oocytes for metaphase II spreads were incubated for 8 minutes instead of 13. Oocytes were subsequently moved into 7μl of fixing solution (1% PFA pH 9.2, 0.15% Triton X-100, 3mM dithiothreitol) on 15-well Multitest Slides (6041505, MP Biomedicals). One oocyte was placed per well, allowing for correct counting of number of chromosomes and chromosome fragments. Oocytes were left to spread overnight. The following day, slides were air-dried in a fume hood and, once fully dried, stored at -80°C.

For immunofluorescence, slides were thawed in PBS for 5 minutes, followed by a 5-minute wash in PBS supplemented with 0.2% BSA, 0.1% Tween 20 (PBT). Then, slides were incubated with blocking solution (10% FBS, 2.5% BSA, 0.1% Tween 20 in PBS) for 1 hour at RT. Primary antibody (CREST serum, a gift from W. Earnshaw, 1:1000 dilution) was diluted in blocking solution and incubated on slides overnight at 4°C. Following 3 x 10-minute washes, slides were incubated with secondary antibody (anti-human Alexa Fluor 488, ThermoFisher, 1:500 dilution), diluted in PBT, for 1.5 hours at RT. The secondary antibody solution also contained 1μg/ml of Hoechst 33342 (62249, ThermoFischer). After 3 x 15-minute washes, slides were mounted in VECTASHIELD Antifade Mounting Medium.

Imaging was performed using a Fluoview FV4000 Confocal Laser Scanning Microscope (Evident Corporation), running cellSens Fluoview software, using a UPLXAPO 40x/1.40 Oil objective on 3X zoom and a SilVIR detector for image acquisition. A small z-stack was captured for each oocyte spread at 0.4μm steps. Z projections were manually assessed for the presence of chromosomal fragments (metaphase II) or broken chromosomes (metaphase I).

### Zygote harvesting and *in vitro* culture

All embryos used in this study were acquired from natural matings. Control and mutant females (non-irradiated and irradiated) were mated with wild-type C57BL/6 males in trios (2 females/1 male) and were checked daily for vaginal plugs. On the day of plug detection (E0.5), females were culled and zygotes were retrieved from oviducts in FHM medium (made in house by the Crick Genetic Modification Services Science Technology Platform, GeMS STP). Zygotes were briefly washed in 1mg/ml Hyaluronidase (H4272, Sigma-Aldrich) solution in warm FHM (<2 minutes total). Afterwards, zygotes were washed thoroughly through 9μl FHM drops under oil, before being washed through 9μl drops of KSOM (made in house by GeMS STP) medium under oil. Zygotes were cultured in KSOM drops for 3 days at 37°C and were assessed daily for development.

### Western blotting

To extract protein samples, pooled, whole ovaries were lysed in 30μl of RIPA buffer (20-188, Millipore). 4-6 ovaries were used per sample, and the whole lysate was loaded in a single gel lane. Protein transfer was performed on PVDF membranes using the iBlot2 system (Invitrogen), at program P0. Blocking was performed in 5% milk solution diluted in Tris-buffered saline, supplemented with 0.1% Tween 20 (TBS-T). Primary antibodies were diluted in blocking solution. Antibodies used: anti-Tubulin (mouse host, 1:3000 dilution, T9026, Sigma-Aldrich), anti-ATR (goat host, 1:50 dilution, sc-21848, Santa Cruz), anti-ATM (rabbit host, 1:100 dilution, 2873T, Cell Signaling Technology). Primary incubation was performed at 4°C O/N. Secondary, HRP-conjugated antibodies were diluted in 5% BSA in TBS-T and were incubated at RT for 90 minutes. Blots were imaged using a ChemiDoc MP imaging system (Bio-Rad).

## References

1. Pepling, M. E. From primordial germ cell to primordial follicle: mammalian female germ cell development. genesis 44, 622–632 (2006).

2. Keeney, S., Giroux, C. N. & Kleckner, N. Meiosis-Specific DNA Double-Strand Breaks Are Catalyzed by Spo11, a Member of a Widely Conserved Protein Family. Cell 88, 375–384 (1997).

3. Bergerat, A. et al. An atypical topoisomerase II from archaea with implications for meiotic recombination. Nature 386, 414–417 (1997).

4. Romanienko, P. J. & Camerini-Otero, R. D. The Mouse *Spo11* Gene Is Required for Meiotic Chromosome Synapsis. Molecular Cell 6, 975–987 (2000).

5. Shinohara, A. & Shinohara, M. Roles of RecA homologues Rad51 and Dmc1 during meiotic recombination. Cytogenet Genome Res 107, 201–207 (2004).

6. Hunter, N. Meiotic Recombination: The Essence of Heredity. Cold Spring Harb Perspect Biol 7, a016618 (2015).

7. Bolcun-Filas, E., Rinaldi, V. D., White, M. E. & Schimenti, J. C. Reversal of female infertility by Chk2 ablation reveals the oocyte DNA damage checkpoint pathway. Science 343, 533–536 (2014).

8. Suh, E.-K. et al. p63 protects the female germ line during meiotic arrest. Nature 444, 624–628 (2006).

9. Schimenti, J. C. Genome integrity checkpoints in mammalian oogenesis. Biol Reprod ioag118 (2026) doi:10.1093/biolre/ioag118.

10. Di Giacomo, M. et al. Distinct DNA-damage-dependent and -independent responses drive the loss of oocytes in recombination-defective mouse mutants. Proceedings of the National Academy of Sciences 102, 737–742 (2005).

11. Ravindranathan, R., Raveendran, K., Papanikos, F., San-Segundo, P. A. & Tóth, A. Chromosomal synapsis defects can trigger oocyte apoptosis without elevating numbers of persistent DNA breaks above wild-type levels. Nucleic Acids Res 50, 5617–5634 (2022).

12. Kogo, H. et al. HORMAD2 is essential for synapsis surveillance during meiotic prophase via the recruitment of ATR activity. Genes Cells 17, 897–912 (2012).

13. Kogo, H. et al. HORMAD1-dependent checkpoint/surveillance mechanism eliminates asynaptic oocytes. Genes to Cells 17, 439–454 (2012).

14. Wojtasz, L. et al. Mouse HORMAD1 and HORMAD2, two conserved meiotic chromosomal proteins, are depleted from synapsed chromosome axes with the help of TRIP13 AAA-ATPase. PLoS Genet 5, e1000702 (2009).

15. Wojtasz, L. et al. Meiotic DNA double-strand breaks and chromosome asynapsis in mice are monitored by distinct HORMAD2-independent and -dependent mechanisms. Genes Dev. 26, 958–973 (2012).

16. Fukuda, T., Daniel, K., Wojtasz, L., Toth, A. & Höög, C. A novel mammalian HORMA domain-containing protein, HORMAD1, preferentially associates with unsynapsed meiotic chromosomes. Exp Cell Res 316, 158–171 (2010).

17. Jiao, X. et al. Aberrant activation of chromosome asynapsis checkpoint triggers oocyte elimination. Nat Commun 16, 2260 (2025).

18. Bai, L. et al. BRCA1 safeguards genome integrity by activating chromosome asynapsis checkpoint to eliminate recombination-defective oocytes. Proc Natl Acad Sci U S A 121, e2401386121 (2024).

19. Qiao, H. et al. Impeding DNA Break Repair Enables Oocyte Quality Control. Molecular Cell 72, 211–221.e3 (2018).

20. Rinaldi, V. D., Bloom, J. C. & Schimenti, J. C. Oocyte Elimination Through DNA Damage Signaling from CHK1/CHK2 to p53 and p63. Genetics 215, 373–378 (2020).

21. Widger, A. et al. ATR is a multifunctional regulator of male mouse meiosis. Nat Commun 9, 2621 (2018).

22. Royo, H. et al. ATR acts stage specifically to regulate multiple aspects of mammalian meiotic silencing. Genes Dev 27, 1484–1494 (2013).

23. Pacheco, S., Marcet-Ortega M., Lange J., Jasin M., Keeney S. & Roig, I. The ATM Signaling Cascade Promotes Recombination-Dependent Pachytene Arrest in Mouse Spermatocytes. PLoS Genet 11, e1005017 (2015).

24. Lange, J. et al. ATM controls meiotic double-strand-break formation. Nature 479, 237– 240 (2011).

25. Gallardo, T., Shirley, L., John, G. B. & Castrillon, D. H. Generation of a germ cell-specific mouse transgenic Cre line, Vasa-Cre. genesis 45, 413–417 (2007).

26. Pan, B. & Li, J. The art of oocyte meiotic arrest regulation. Reprod Biol Endocrinol 17, 8 (2019).

27. Pacheco, S., Maldonado-Linares, A., Garcia-Caldés, M. & Roig, I. ATR function is indispensable to allow proper mammalian follicle development. Chromosoma 128, 489–500 (2019).

28. Plug, A. W. et al. ATM and RPA in meiotic chromosome synapsis and recombination. Nat Genet 17, 457–461 (1997).

29. Baudat, F., Manova, K., Yuen, J. P., Jasin, M. & Keeney, S. Chromosome Synapsis Defects and Sexually Dimorphic Meiotic Progression in Mice Lacking Spo11. Molecular Cell 6, 989–998 (2000).

30. Mahadevaiah, S. K. et al. Recombinational DNA double-strand breaks in mice precede synapsis. Nat Genet 27, 271–276 (2001).

31. Pittman, D. L. et al. Meiotic prophase arrest with failure of chromosome synapsis in mice deficient for Dmc1, a germline-specific RecA homolog. Mol Cell 1, 697–705 (1998).

32. Yoshida, K. et al. The Mouse RecA-like Gene Dmc1 Is Required for Homologous Chromosome Synapsis during Meiosis. Molecular Cell 1, 707–718 (1998).

33. Barlow, C. et al. Atm deficiency results in severe meiotic disruption as early as leptonema of prophase I. Development 125, 4007–4017 (1998).

34. Barchi, M. et al. Surveillance of different recombination defects in mouse spermatocytes yields distinct responses despite elimination at an identical developmental stage. Mol Cell Biol 25, 7203–7215 (2005).

35. Bakkenist, C. J. & Kastan, M. B. DNA damage activates ATM through intermolecular autophosphorylation and dimer dissociation. Nature 421, 499–506 (2003).

36. Burrows, A. E. & Elledge, S. J. How ATR turns on: TopBP1 goes on ATRIP with ATR. Genes Dev. 22, 1416–1421 (2008).

37. Zou, L. & Elledge, S. J. Sensing DNA damage through ATRIP recognition of RPA-ssDNA complexes. Science 300, 1542–1548 (2003).

38. Namiki, Y. & Zou, L. ATRIP associates with replication protein A-coated ssDNA through multiple interactions. Proceedings of the National Academy of Sciences 103, 580–585 (2006).

39. Choi, J.-H. et al. Reconstitution of RPA-covered single-stranded DNA-activated ATR-Chk1 signaling. Proceedings of the National Academy of Sciences 107, 13660–13665 (2010).

40. ElInati, E. et al. The BCL-2 pathway preserves mammalian genome integrity by eliminating recombination-defective oocytes. Nat Commun 11, 2598 (2020).

41. Cai, Z., Chehab, N. H. & Pavletich, N. P. Structure and Activation Mechanism of the CHK2 DNA Damage Checkpoint Kinase. Molecular Cell 35, 818–829 (2009).

42. Matsuoka, S. et al. Ataxia telangiectasia-mutated phosphorylates Chk2 in vivo and in vitro. Proc Natl Acad Sci U S A 97, 10389–10394 (2000).

43. Chehab, N. H., Malikzay, A., Appel, M. & Halazonetis, T. D. Chk2/hCds1 functions as a DNA damage checkpoint in G1 by stabilizing p53. Genes Dev. 14, 278–288 (2000).

44. Hirao, A. et al. DNA damage-induced activation of p53 by the checkpoint kinase Chk2. Science 287, 1824–1827 (2000).

45. Matsui, Y., Nagano, R. & Obinata, M. Apoptosis of fetal testicular cells is regulated by both p53-dependent and independent mechanisms. Mol. Reprod. Dev. 55, 399–405 (2000).

46. Yamada, S. et al. Molecular structures and mechanisms of DNA break processing in mouse meiosis. Genes Dev. 34, 806–818 (2020).

47. Huang, P. et al. Molecular Mechanisms of Parthanatos and Its Role in Diverse Diseases. Int J Mol Sci 23, 7292 (2022).

48. Zha, S., Sekiguchi, J., Brush, J. W., Bassing, C. H. & Alt, F. W. Complementary functions of ATM and H2AX in development and suppression of genomic instability. Proc Natl Acad Sci U S A 105, 9302–9306 (2008).

49. Brown, E. J. & Baltimore, D. ATR disruption leads to chromosomal fragmentation and early embryonic lethality. Genes Dev 14, 397–402 (2000).

50. Ruzankina, Y. et al. Deletion of the developmentally essential gene ATR in adult mice leads to age-related phenotypes and stem cell loss. Cell Stem Cell 1, 113–126 (2007).

51. Lin, Z. et al. Mettl3-/Mettl14-mediated mRNA N6-methyladenosine modulates murine spermatogenesis. Cell Res 27, 1216–1230 (2017).

52. Bosma, G. C., Custer, R. P. & Bosma, M. J. A severe combined immunodeficiency mutation in the mouse. Nature 301, 527–530 (1983).

53. McKey, J., Cameron, L. A., Lewis, D., Batchvarov, I. S. & Capel, B. Combined iDISCO and CUBIC tissue clearing and lightsheet microscopy for in toto analysis of the adult mouse ovary†. Biol Reprod 102, 1080–1089 (2020).

54. Mineo, A. et al. The sex and reproductive plasticity of intestinal muscles instruct gut size. Cell 189, 6078–6096.e9 (2026).

